# The Emergence of Egalitarianism in a Model of Early Human Societies

**DOI:** 10.1101/166116

**Authors:** Guillaume Calmettes, James N. Weiss

## Abstract

How did egalitarianism emerge in early human societies? In contrast to dominance hierarchies in non-human primates, human simple forager bands are typically egalitarian, with male hunters often serving as the collective alpha. Here we present a thermodynamics-inspired simple population model, based on stochastic optimization of dominance relationships, in which a dominance hierarchy of individuals with exclusively self-centered characteristics (the desire to dominate, resentment at being dominated) transitions spontaneously to egalitarianism as their capacity for language develops. Language, specifically gossip, allows resentment against being dominated to promote the formation of antidominance coalitions which destabilize the alpha position for individuals, leading to a phase transition in which a coalition of the full population suddenly becomes dominant. Thus, egalitarianism emerges suddenly as the optimal power sharing arrangement in a population of selfish individuals without any inherently altruistic qualities. We speculate that egalitarianism driven by punishment for exhibiting alpha-like behavior may then set the stage for genuinely altruistic traits to propagate as predicted by game theory models. Based on model simulations, we also predict that egalitarianism is a pre-condition for adaptation of tools as weapons. Potential implications for origins of human moral belief systems are discussed.

## Introduction

Non-human primates generally form hierarchal social communities with either dominant individuals (alphas) or small coalitions at the top and lower-ranking individuals below [1]. In contrast, extant small-scale human simple forager bands of mobile hunter-gatherers, who arguably resemble the earliest human communities in the late Pleistocene era [1, 2], typically exhibit egalitarian power-sharing among male hunters, who make major decisions by consensus [3, 4]. In these egalitarian anti-dominance coalitions, political power is shared equitably, with a variety of cultural status leveling mechanisms (ridicule, ostracism, shunning, exile and execution) discouraging individuals from adopting alpha-like behavior. In effect, a coalition of the weak enforces a moral code that keeps aggressive upstarts from selfishly dominating the group to monopolize reproductive and other resources. These egalitarian simple forager bands typically range in size from 20-30 related and unrelated individuals, although they also form larger networks of bands. After bands coalesce into tribes, chiefdoms and states with hundreds or thousands of members, however, such communal decision-making becomes increasingly unwieldy, and dominance hierarchies commonly reappear in the form of big men, chiefs, kings, dictators and political leaders [3, 4].

Multiple socio-ecological factors have been proposed to play a role in shaping the transition from hierarchal social structure in non-human primates to egalitarianism in human simple foragers (the egalitarian transition), such as the need for cooperation to hunt large game with weapons, to gain the nutritional advantages of meat-sharing between infrequent kills, to wage warfare effectively against competing human communities in relation to the abundance and defensibility of contested resources, variance in male quality and to develop language and cognitive skills required for cooperative behavior [5-12]. Together, these socio-ecological factors created a “socio-cognitive niche” that distinguished early humans from other primates [13]. However, which of these multiple factors played the major role in initiating the egalitarian transition, as opposed to synergistically reinforcing it once it had occurred, is unclear. Some factors, notably large game hunting with weapons, would appear to require high levels of cooperation as a precondition. Moreover, in extant simple foragers, egalitarianism has been observed across a diverse range of socio-ecological conditions, from tropical rain forests to deserts and the arctic, suggesting that a universal factor, rather than purely local environmental conditions, may be at play.

By facilitating communication of third party information, complex language development has been proposed as a critical factor allowing large-scale coalitions to form in early human communities [8, 14, 15]. Also, a recent stochastic model, in which individuals in a population were randomly assigned affinities to assist (or not assist) other individuals during a conflict, showed that egalitarianism could arise suddenly as a phase transition when the average affinity strength reached a critical value [16, 17]. More generally, recent applications of statistical physics to game theory models of human cooperation have shown that both continuous and discontinuous (first and second order) phase transitions occur commonly when punishment and/or reward elements are incorporated into public goods games (for reviews see [18, 19]. Here we synthesize these ideas into a novel thermodynamics-inspired population model based on stochastic optimization to illustrate how complex language development, in the form of gossip, could act as the universal factor triggering the egalitarian transition, analogous to a thermodynamic phase transition. The model population consists of individuals with no inherently altruistic attributes, just self-centered attributes, namely an aggressive will to dominate and a bitter resentment against being dominated. We show that when language skills reach a critical level at which gossip becomes the primary means of reinforcing social ties, the egalitarian transition occurs spontaneously as a phase transition. At this critical point, individuals who resent being dominated become capable of forming and sustaining coalitions that make the individual alpha position progressively unstable, ultimately motivating its avoidance by all members of the population due to retaliation (or fear of retaliation) by a stronger anti-dominance coalition. The implications of this model for adaptation of tools as weapons and the origins of moral belief systems are discussed.

## Results

### Model of primate society

Our model population emulates the size of a typical human male hunter-dominated simple forager band of 30, half of whom are males who become eligible for the dominant male alpha position as they mature into their prime age range. Each individual is characterized by a parameter αβ, which contains two self-centered components, aggressiveness and bitterness. The aggressiveness component (α) quantifies the will and physical ability to dominate other individuals in the population in order to monopolize reproductive and other resources, reflecting the motivation/ability to achieve as high a dominance rank as possible. The bitterness component (β) quantifies resentment at being dominated by other individuals, driving the motivation to avoid a low dominance rank. In most simulations, values of αβ were randomly assigned from a truncated exponential distribution centered at 0.10 (Fig. 1A). In this case, the values of αβ ranged from 1-10 times less than the median in 50% of the population, and 1-10 times greater than the median in the other half. In some simulations, we used a truncated Gaussian distribution (mean 0.50, standard deviation 0.12) or uniform distributions from [0.01, 1.00] for comparison (Fig. 1A). In these cases, the values of αβ ranged from 1-50 times less (i.e. from 0.01 to 0.5) than the median in 50% of the population, and 1-2 times greater (i.e. from 0.5 to 1.0) than the median in the other half. We assumed that the ability to dominate other individuals is zero at birth, rises to a peak and then declines to zero at death. To incorporate this age-dependence into the dominance rank, we randomly assigned an initial age a_0_ (from 0 to 50 years) to each individual in the population. The dominance rank of each individual over time was then determined by their individual Dominance Score (DS), defined as DS = *αβ* * *δ*(*t*), where *δ*(*t*) = sin[*π*((a_0_ + *t*) *mod*50)/50]. Thus, using this half sine wave function for, the DS was zero at birth, rose smoothly to a maximum of *αβ* at age 25 years, and then declined smoothly to zero again at the time of death at 50 years of age (as illustrated in Fig. 2A). This ensured that neither children nor elderly individuals could attain the dominant alpha position (i.e. the highest DS). Individuals died at age 50 and were replaced by new individuals at age 0. Thus, at any given point in time, the appropriately middle-aged individual with the strongest will and physical ability to dominate, coupled with the greatest resentment against being dominated, had the highest DS and therefore assumed the top rank, and the individual with the lowest score assumed the bottom rank. Fig. 2A shows how the DS of 5 individuals (and their replacements upon reaching age 50) changed over 100 years, illustrating a dominance hierarchy in which the dominant alpha position shifted intermittently to different individuals with higher individual DS (color shaded areas) when they reached their prime age range. In most simulations, individuals who died at age 50 were replaced with new individuals at age 0 with randomly selected new values of αβ, reflecting no direct genetic inheritance between the dead individuals and their replacements. This simulates the situation in which the replacements represent new births from different parents at the time that a non-kin individual dies. This is analogous to a randomly changing gene pool controlling the aggressiveness-bitterness factor αβ in the population without any directed selective pressure, ensuring that the egalitarian transition in the model is not related to selection for aggressiveness-bitterness traits. For comparison, in some simulations we randomly assigned the same αβ values to the new individuals, as if they genetically inherited 100% of the characteristics (genes) of the individuals whom they replaced, equivalent to an unchanging gene pool.

**Fig. 1.**
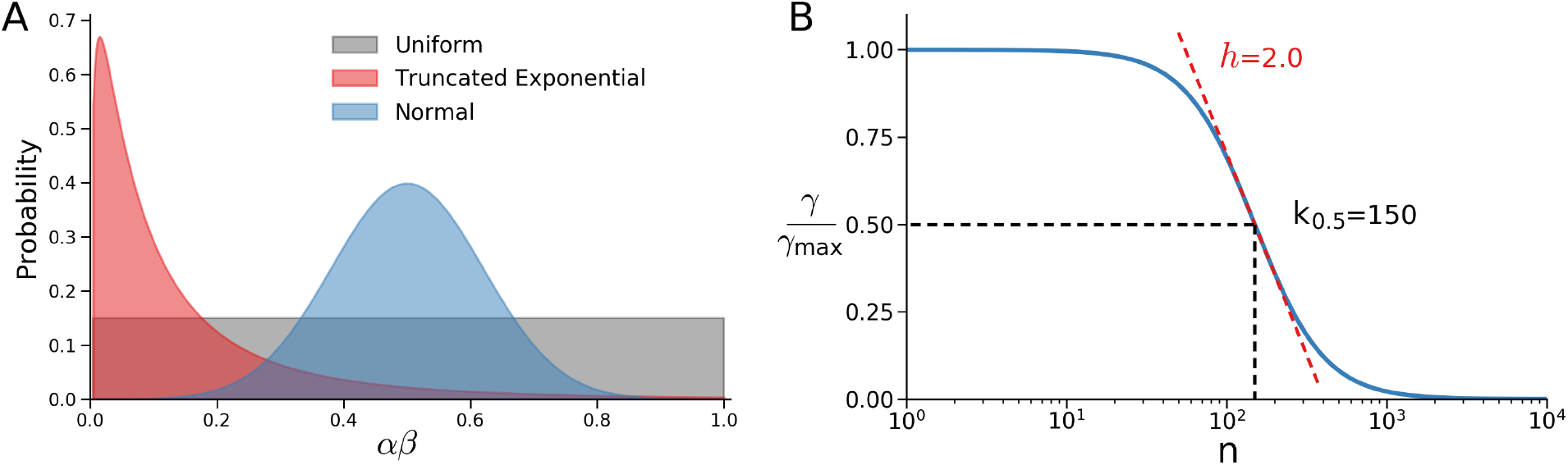
**A.** Exponential (red), normal (blue) and uniform (gray) distributions used to randomly select values of the aggressiveness-bitterness factor αβ. **B.** Hill equation formulation used to compute the gossip factor γ as a function of coalition size (n), where γ_max_ is the maximum value of γ, K_0.5_ is coalition size at which γ/γ_max_=0.5 and *h* is the Hill coefficient.

**Fig. 2.**
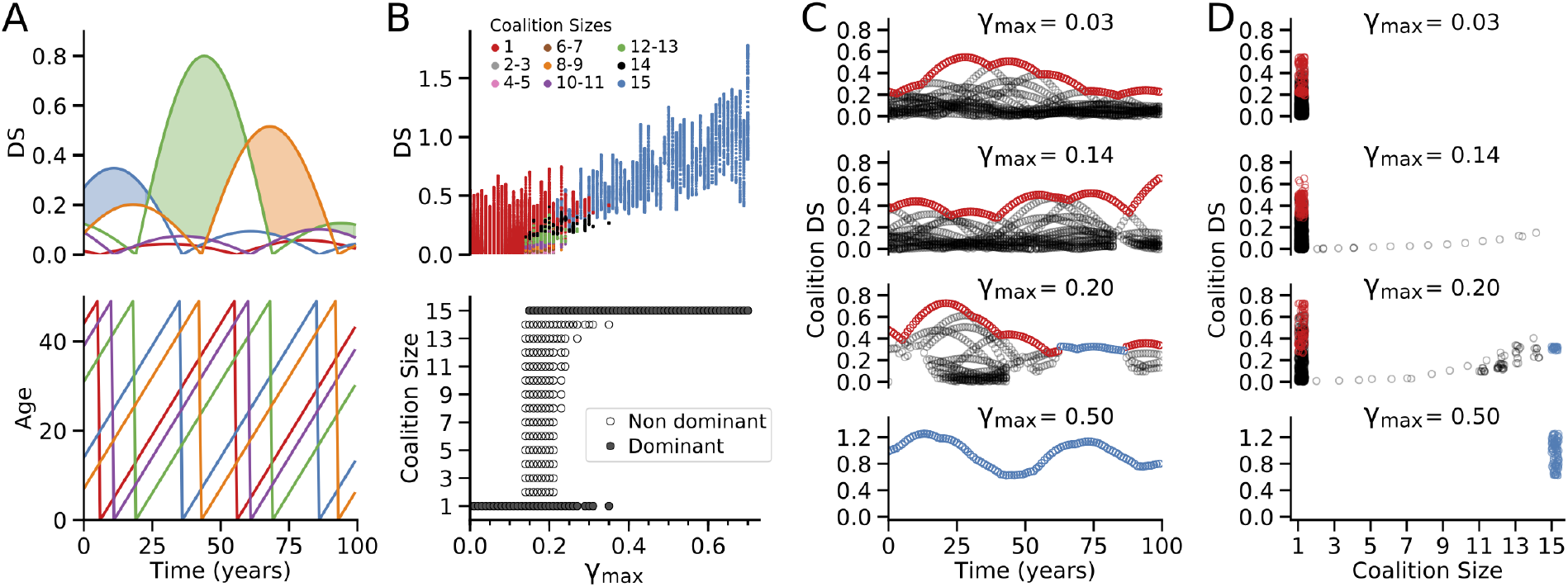
Transition from alpha dominance to egalitarianism. **A.** Dominance Scores (DS) for 5 representative individuals *(upper panel)* with aging *(lower panel)*. Shading indicates the dominant individual alpha with the highest DS. **B.** *Upper panel:* DS of individuals (red), nondominant coalitions of various sizes (color-coded) and the egalitarian coalition of the full population (blue) as γ_max_ increases, for a population of 15 individuals with K_0.5_=150 and *h*=2.0. For each γ_max_ value, the DS values shown span 100 years. *Lower panel:* Effect of increasing γ_max_ on the dominant coalition size (closed symbols), showing a transition from a dominant alpha individual to the egalitarian coalition of all 15 individuals at γ_max_=0.13. From γ_max_=0.13-0.38, the population intermittently switches between a dominant alpha individual and the egalitarian coalition as DS values change with age. During the periods when the alpha dominates, nondominant coalitions can form (open symbols). **C.** Expanded traces of DS values over 100 years for 4 values of γ_max_ (0.03, 0.14, 0.20 and 0.50). Non-dominant individuals and coalitions are shown in black, dominant individuals in red, and dominant egalitarian coalitions in blue. **D.** Corresponding individual and coalition DS values for different coalition sizes in D.

To incorporate language development, we introduced a global variable, the gossip factor γ, whose value reflects the state of language skills (or, equivalently, genes controlling language skills) that have evolved in the population. We do not attempt to explain how or why language evolved, merely that it did and eventually replaced grooming as the major mechanism for reinforcing social bonds. Based on previous arguments that language played a critical role in large scale coalition formation in early humans [8, 14, 15], we postulate that through gossip, individuals gain the ability to communicate with third parties to share their resentment against being dominated by higher ranking individuals, including the alpha, and to formulate a proactive plan to come to each others* aid in the event of a conflict. Thus, gossip serves as the primary vehicle for forming small or large coalitions. However, since gossiping to share information and reinforce social bonding in coalitions consumes time that could otherwise be spent for alternative vital activities necessary for survival, we assume that the cost of gossip at forming and maintaining coalitions increases as coalition size *N* increases, which we modeled as a Hill function (Fig. 1B):

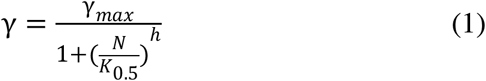

where γ_max_ is the maximum value of the gossip variable γ, K_0.5_ is the coalition size *N* at which γ=0.5*γ_max_, and *h* is the Hill coefficient. For γ_max_>0, the ability to communicate privately to third parties (i.e. to gossip) allows like-minded individuals, who resent being dominated by another individual or coalition, to form their own coalition whose collective DS is determined by the sum of the individual DS of all coalition members multiplied by γ(*N*), i.e.

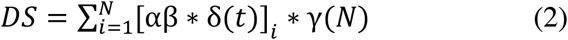

In the simulations, individuals were randomly selected to form coalitions that increased their collective DS and thereby improve their dominance rank. Specifically, individuals were permitted to join a coalition <u>if and only if</u> their inclusion increased both the coalition's collective DS and was greater than the individual's DS (i.e. all coalition members improved their dominance rank when a coalition formed). The sociopolitical standing in the population was then ranked in the order of the highest to lowest DS of both individuals and coalitions.

### Transition from alpha hierarchy to egalitarianism

With this model, we used γ_max_ as the control parameter to study how the development of complex language skills allowed resentful individuals to form gossip-driven coalitions in order to improve their sociopolitical ranking. The value of K_0.5_ in Eq.(1) was set at 150, a rough estimate of the maximum number of modern humans for whom it is practical to gossip regularly enough to reinforce the social ties and trust levels necessary to maintain a cohesive coalition [20]. The value of the Hill coefficient *h* was arbitrarily set at 2.0 (see later for analysis of the effects of *h*). Individual and coalition DS values were computed as the gossip parameter γ_max_ was increased from zero (no gossip skills) to positive values (progressively stronger gossip skills) in 0.01 increments as follows: 1) For each γ_max_ value, individuals (*N=*1) or coalitions (*N*=2-15) were ranked from lowest to highest DS. Individuals from the lowest ranking coalition were then allowed to defect one-by-one to randomly join another individual or coalition as long as the new coalition's collective DS increased and was greater than the individual's DS. This process was iterated until no further defections caused any coalition's DS to increase further, i.e. an equilibrium was reached corresponding to an optimized stable sociopolitical power structure. 2) Time was then advanced by 1 year, and the algorithm repeated. 3) This process was iterated 100 times (100 years) for each γ_max_ value.

The results from a typical simulation are shown in Fig. 2B-D, all using the same initial set of randomized values of αβ and *a*_*o*_. For γ_max_ values <0.15, individuals always had a higher DS than any hypothetical coalition (upper panel), so that the highest-ranking individual remained the dominant alpha with lower-ranking individuals forming a pecking order (Fig. 2B, lower panel). When γ_max_ reached 0.15, however, a coalition of the full population (*N*=15) intermittently achieved a DS that exceeded the highest ranking individual's DS, thereby switching abruptly from an alpha hierarchy with a single dominant individual (red dots) to an egalitarian coalition of all 15 individuals (blue dots). However, the population could still revert back to an alpha hierarchy intermittently as individuals naturally endowed with a very high αβ reached prime age such that their individual DS exceeded that of the egalitarian coalition. During these switches, non-dominant beneficial coalitions of smaller size formed (*N*=2-14, other colored dots in the top panel, open circles in lower panel), which improved the status of their members (i.e. a higher coalition DS compared to individual DS) even though the population was still dominated by an individual alpha. For γ_max_ > 0.35, the population stabilized to full egalitarianism without intermittent reversions to a dominance hierarchy. The smaller coalitions also disappeared, since DS of the egalitarian coalition was higher than that of any individual or smaller coalition. Thus, the population was always dominated by either an alpha individual or an egalitarian coalition of the full population, and never by a smaller coalition for these parameter settings (K_0.5_=150 and *h*=2.0, see later), resembling a sudden phase transition rather than a gradual process in which small co-dominant coalitions gradually became progressively larger and finally egalitarian. Fig. 2C shows the evolution of DS values over a 100 year period for four values of γ_max_. γ_max_=0.03 produced a dominant individual alpha (red dots) without co-existing non-dominant coalitions (black dots). γ_max_=0.14 produced a dominant individual alpha (red dots) with co-existing nondominant coalitions (black dots). γ_max_=0.20 produced a dominant individual alpha (red dots) intermittently switching to a dominant egalitarian coalition of the full population (blue dots). γ_max_=0.50 produced a permanently dominant egalitarian coalition. The corresponding range of DS of individuals and coalitions of various sizes for these different γ_max_ values are shown in Fig. 2D, with all the dominant individuals/coalitions during this 100 year period indicated in red for dominant alpha individuals and in blue for dominant coalitions.

In the simulation in Fig. 2, new individuals replacing dead individuals were assigned new randomly selected values of αβ, emulating the case in which the gene pool controlling these factors fluctuates randomly over time. To ensure that these random fluctuations in αβ were not responsible for driving the egalitarian transition, we also performed simulations in which new individuals were assigned the identical αβ values to those whom they replaced to emulate a stable αβ gene pool. The results were similar, except that the intergenerational dominance pattern repeated itself cyclically every 50 years. The type of distribution from which the αβ values were randomly assigned also had no major effects, since replacing the truncated exponential distribution of αβ with either the truncated Gaussian or uniform distributions described above yielded qualitatively similar results.

### Model robustness model in relation to free-riders

We assumed in the model that the coalition DS computed from Eq.(2) accurately determined the dominance rank of the coalition in the population, i.e. that coalition members always cooperated fully to realize the full potential of the computed coalition DS. In reality, however, sustained cooperation by a coalition is vulnerable to free-riders, i.e. individuals who join a coalition to share the advantages of its dominance rank, but then cheat by not contributing fully when conflicts with bullies or competing coalitions arise. Because of the free-rider problem, the origins of large scale cooperation in the evolution of human societies is highly debated, since within-group competition favors selfish over altruistic genes in game theoretic models [21-24]. In our model, free-riders would decrease a coalition's effective DS (i.e. increase the cost and decrease the benefit of being a member of the coalition). To examine our model's susceptibility to free-riders, we modified Eq. 2 to include an additional factor, the laziness factor λ, to represent free-rider contamination:

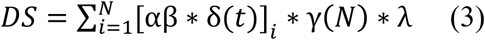

The laziness factor λ, whose value ranges from 0 to 1, is an aggregate parameter collectively reflecting the personalities/compatibilities/free-riding tendencies of individuals in a potential coalition in terms of their likelihood to cheat, deceive, lose nerve in a conflict, etc. Thus, λ=0 represents an ineffective coalition consisting of all uncooperative free-riders (equivalent to high cost, no benefit), λ=1 represents a maximally effective coalition of all cooperative individuals (equivalent to low cost, high benefit), and intermediate values represent mixtures. Each potential new coalition during the course of the year was assigned a value of λ randomly selected from a uniform distribution between [0,1]. A coalition was only allowed to form if its DS after multiplication by its randomly selected λ value met the same criteria described earlier, namely that the new member increased the coalition's collective DS and exceeded the individual DS values of all its constituent members. With the laziness factor λ included, the population still underwent a similar egalitarian transition (Fig. 3A), although the number of required iterations increased modestly and the transition occurred at a slightly higher value of the gossip parameter γ_max_ (0.18 vs 0.15 for the simulation without λ in Fig. 2B). The randomly assigned laziness factor γ value of the final egalitarian coalition was always >0.95. Similar results were obtained using a truncated Gaussian distribution centered at 0.5 (95% CI [0.1, 0.9]) to randomly select the value of the laziness factor λ. In the above simulations, we assumed that a new coalition with a higher DS displaced lower-ranking coalitions with 100% certainty. However, this assumption was also not critical. If the probability of a new coalition displacing a lower-ranking coalition was reduced to 50%, the model still achieved an optimal stable configuration in which no further defections improved any individual's or coalition's DS, although the number of required iterations again increased modestly (Fig. 3B).

**Fig. 3.**
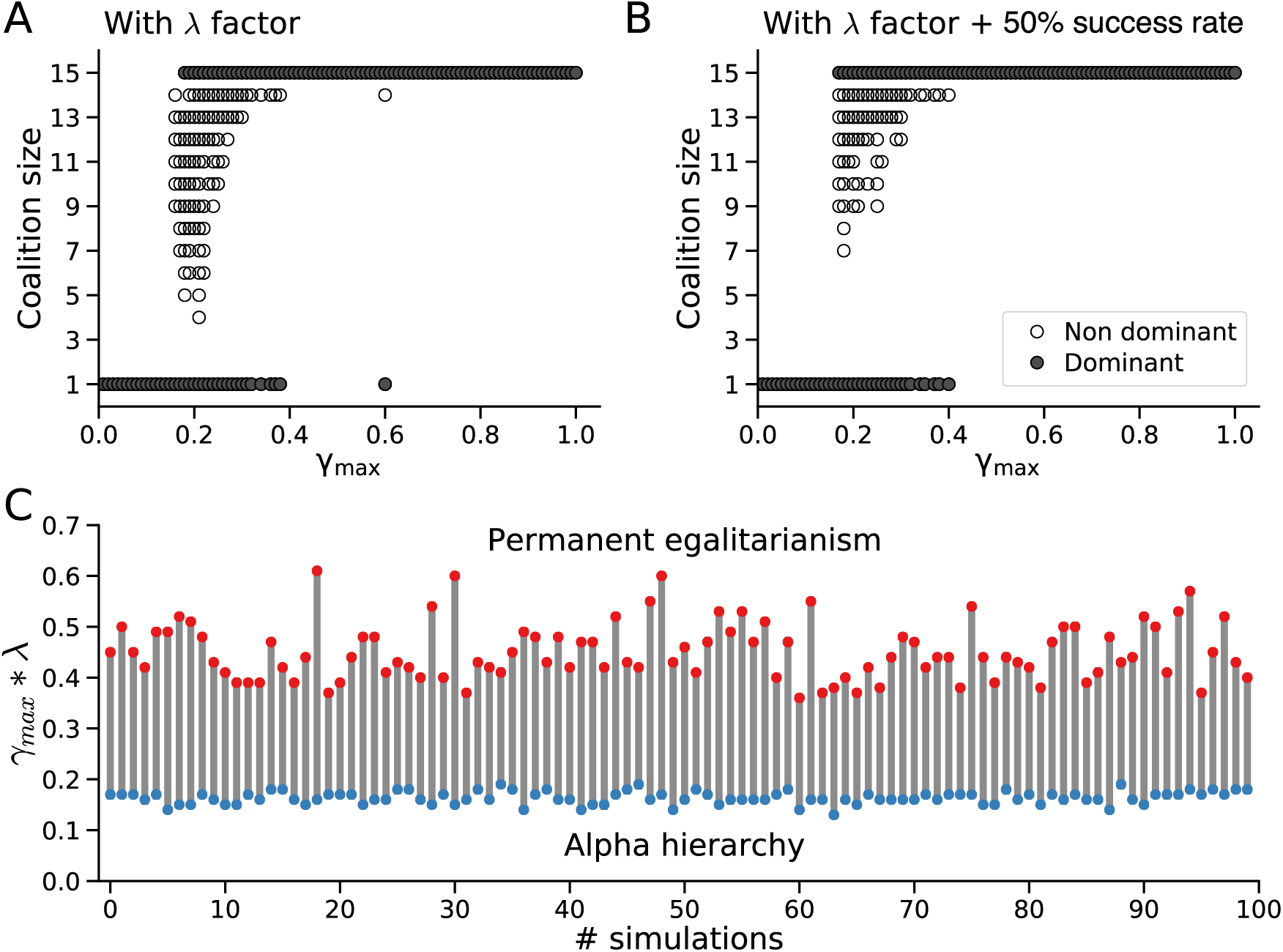
Effects of free-riders and reduced success rate. **A.** Similar to the lower panel in Fig. 2B, when the laziness factor λ in Eq.(3) is added to the model to simulate free-riders (see text), the hierarchy to egalitarian transition still occurs, although at a slightly higher γ_max_ value. Closed symbols show the dominant coalition size during a 100 years period for each γ_max_; open symbols show non-dominant coalitions when an alpha individual has the top rank. **B.** The same conditions incorporating the λ factor in the model, but also with the success rateof a coalition with a higher DS replacing a lower-ranking coalition or individual reduced to 50%. **C.** Predicted values of γ_max_•λfrom Eq.(4) at which the population transitions from alpha hierarchy to intermittent egalitarianism (blue symbols) to permanent egalitarianism (red symbols), compared to the numerical simulation results (gray lines) for 100 randomized trials of different initial age distributions.

Thus, the egalitarian transition in this model is robust to the free-rider problem, and is also generally robust as can be readily deduced from Eq.(1) and Eq.(3). Since the gossip factor g is directly proportional to γ_max_ in Eq.(1), and the coalition DS in Eq.(3) is also directly proportional to g, then a large enough value of γ_max_ will inevitably cause the coalition DS 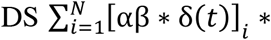 λ*γ to exceed the largest individual DS in the population [αβ *δ*(*t*)]_max_ at any given time point *t*. Analytically, the γ_max_ value at which the population transitions from hierarchical to egalitarian for a given randomly selected value of λ can thus be algebraically derived from Eqs.(1) and (3) as:

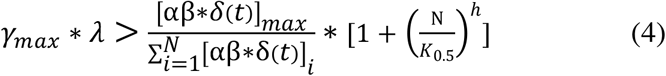

For a time interval Dt, the population will be hierarchical if the condition in Eq.(4) is never met, intermittently egalitarian if Eq.(4) is met at some time points during Δt, and permanently egalitarian if Eq.(4) is always met during Δt. We validated this analytical solution using the numerical simulation conditions in Fig. 3C, for which *N*=15, K_0.5_=150, *h=*2. For 100 random assignments of initial ages *a*_0_ to the population, the predicted *γ_max_ * λ* values at which the population transitioned from hierarchical to intermittently egalitarian to permanently egalitarian over a 100 year time interval calculated from Eq.(4) showed excellent agreement with those obtained directly from the numerical simulations (Fig. 3C). This demonstrates that transitions from alpha hierarchy to intermittency to permanent egalitarianism can be analytically predicted from global statistical macroscopic parameters of the population in Eq.(4) without knowledge of the each individual's microscopic details, similar to the mean field assumption in a thermodynamical process.

### Non-dominant coalitions

The transition from a dominant alpha individual to an egalitarian antidominance coalition as the gossip parameter γ_max_ increased reflects the self-serving ability of individuals to improve their dominance rank by forming coalitions as the third party communicative power of gossip increases. In the simulations in Fig. 2, the algorithm reported only the maximal DS that each individual achieved as a member of a coalition, ignoring less powerful coalitions that individuals may have joined along the way. However, whether global power is controlled by a dominant alpha individual or an egalitarian coalition, it can still be advantageous for individuals to form local non-dominant coalitions that improve their DS so that they can exert hierarchical influence over other non-dominant coalitions or individuals. In this case, any unresolved conflicts between competing non-dominant coalitions and/or individuals would ultimately be settled by the dominant alpha individual or dominant egalitarian coalition depending on the range of gossip parameter γ_max_. To analyze the influence of the gossip parameter γ_max_ on the formation of beneficial non-dominant coalitions in the model, we computed all possible combinations of individuals that improved their collective DS when they formed coalitions of any size (Fig. 4A, black trace). We also computed all possible coalitions that improved the DS of the highest-ranking individual (Fig. 4A, blue trace). For low values of the gossip parameter γ_max_, there were no beneficial coalitions of any size, but when γ_max_ reached 0.13 in this simulation, the number of beneficial coalitions for all individuals increased steeply (inset, Fig. 4A). In contrast, there were no beneficial coalitions for the highest-ranking individual until the gossip parameter γ_max_ reached 0.18. At γ_max_=0.18, the population switched abruptly to an egalitarian coalition, and the number of non-dominant beneficial coalitions continued to increase, reaching a maximum at γ_max_=0.6. Fig. 4B shows the distribution of non-dominant coalition sizes at two values of the gossip parameter γ_max_ (0.15 and 0.25), both for all individuals (upper panels) and for the highest-ranking individual (lower panels) in this simulation.

**Fig. 4.**
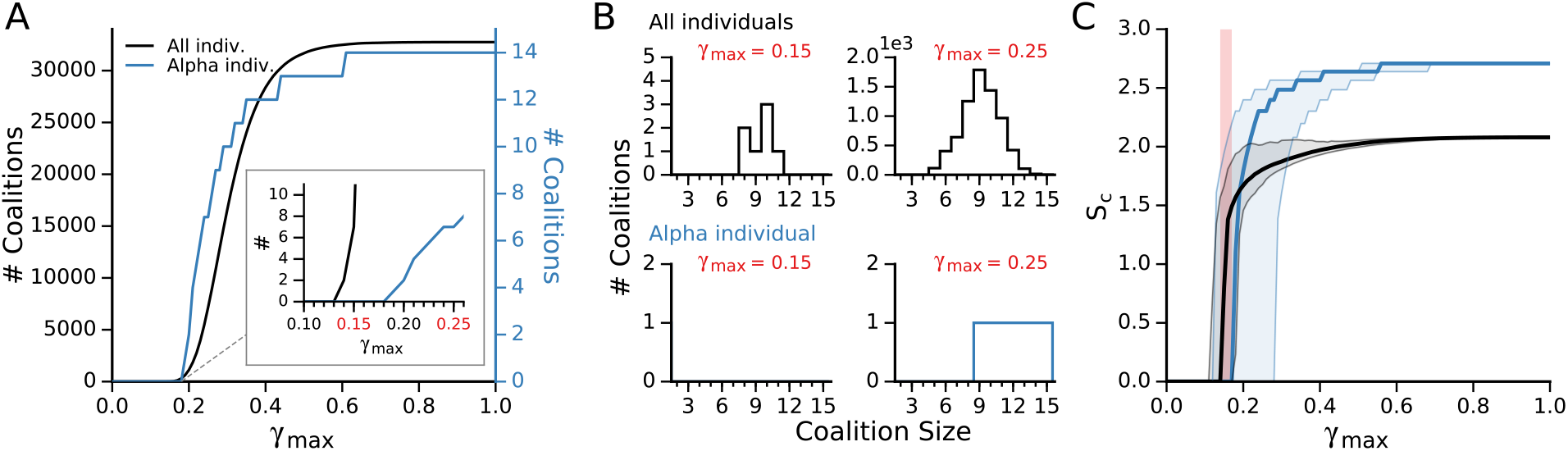
Non-dominant coalitions. **A.**Number of possible beneficial coalitions improving the DS of all individuals (black) or the highest-ranking individual (blue) as γ_max_ increased, for a population of 15 individuals with initially randomized α and β values, K_0.5_=150? and *h*=2.0. Inset shows that beneficial coalitions for all individuals start at a lower γ_max_ compared to the highest-ranking individual. B. Corresponding distributions of beneficial coalition sizes for all individuals (upper graphs) and the highest-ranking individual (lower graphs) at the two different values of γ_max_ in A. C. Coalition entropy (*S*_*c*_) calculated for 1,000 different initial randomizations of α and β, showing the median (thick lines) and 95% confidence intervals (shading). Black curves are for all individuals, blue curves for the highest-ranking individual. Pink shading shows the region in which beneficial coalitions coexist with a dominant alpha individual.

Fig. 4A represents the beneficial non-dominant coalitions for a single initial randomization of αβ values assigned to the population. To analyze the statistical properties of forming beneficial coalitions, we computed the information entropy for 1,000 sets of random initial values of αβ (Fig. 4C). We defined the coalition entropy *S*_*c*_ in standard fashion from the probability distribution *P(ω)* that a randomly-selected individual could improve its DS by joining a coalition as:

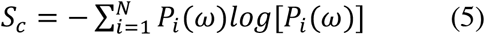

*S*_*c*_ remained low until a critical value of the gossip parameter γ_max_ was reached (median value 0.15), at which point *S*_*c*_ increased steeply (black curve) corresponding to the onset of many possible beneficial coalitions (equivalent to entropic microstates) within a dominance hierarchy (pink shading). The coalition entropy for the highest ranking individual, on the other hand, remained zero until the gossip parameter γ_max_ reached a median value of 0.17 (blue curve), at which point the highest ranking individual joined a coalition signaling the transition to egalitarianism. The black and blue shading indicates the 95% confidence intervals computed from 1,000 different simulations for each curve.

We interpret these findings to indicate that as the gossip parameter γ_max_ increases, manybeneficial coalitions of various sizes become possible which will increase the DS of an individual by joining that coalition, even before the population becomes fully egalitarian. This is compatible with a society of nested, variably sized coalitions exerting local hierarchical influence over each other, whether global power is controlled by a dominant alpha individual or an egalitarian coalition of the full population. The implicit benefit in joining such a coalition is the increased dominance rank achieved through an improved DS, whereas the implicit cost is sharing risk of injury during conflicts and power with other coalition members.

### Effects of the K_0.5_ and Hill coefficient h

The K_0.5_ value in this model has an important role in determining the dominant coalition size in the population. If the Hill relationship (Fig. 1B) is removed from Eq.(1), such that γ=γ_max_, the dominant coalition is always the full population when the gossip parameter γ_max_ reaches its threshold value. Similarly, when K_0.5_ is much greater than the total population size (such as K_0.5_=150 and a maximum coalition size *N* of the full population of 15 in Figs. 2-4), the population always transitions suddenly from dominance hierarchy to full egalitarianism at the critical γ_max_ value. When the K_0.5_ value is smaller than the total population size, however, this is not the case. This is illustrated in Fig. 5A & B, for γ_max_=1.0 (well above the threshold for the egalitarian transition in Fig. 2) in a total population of 15 individuals. The K_0.5_ value was initially set at 1 and increased by 1 every 100 years. As the K_0.5_ value increased (Fig. 5A, black trace), the dominant coalition size increased progressively as smaller coalitions disappeared due to defections to coalitions with higher DS (Fig. 5B), consolidating the population into approximately 15/K_0.5_ coalitions (Fig. 5A, blue trace). As the K_0.5_ value approached 15, the highest DS shifted to the full population of 15 individuals (Fig. 5B), creating a fully egalitarian society at a value of K_0.5_ well below the conventionally estimated maximum number of 150 modern humans for whom the power of gossip is capable of maintaining an effective coalition [20].

**Fig. 5.**
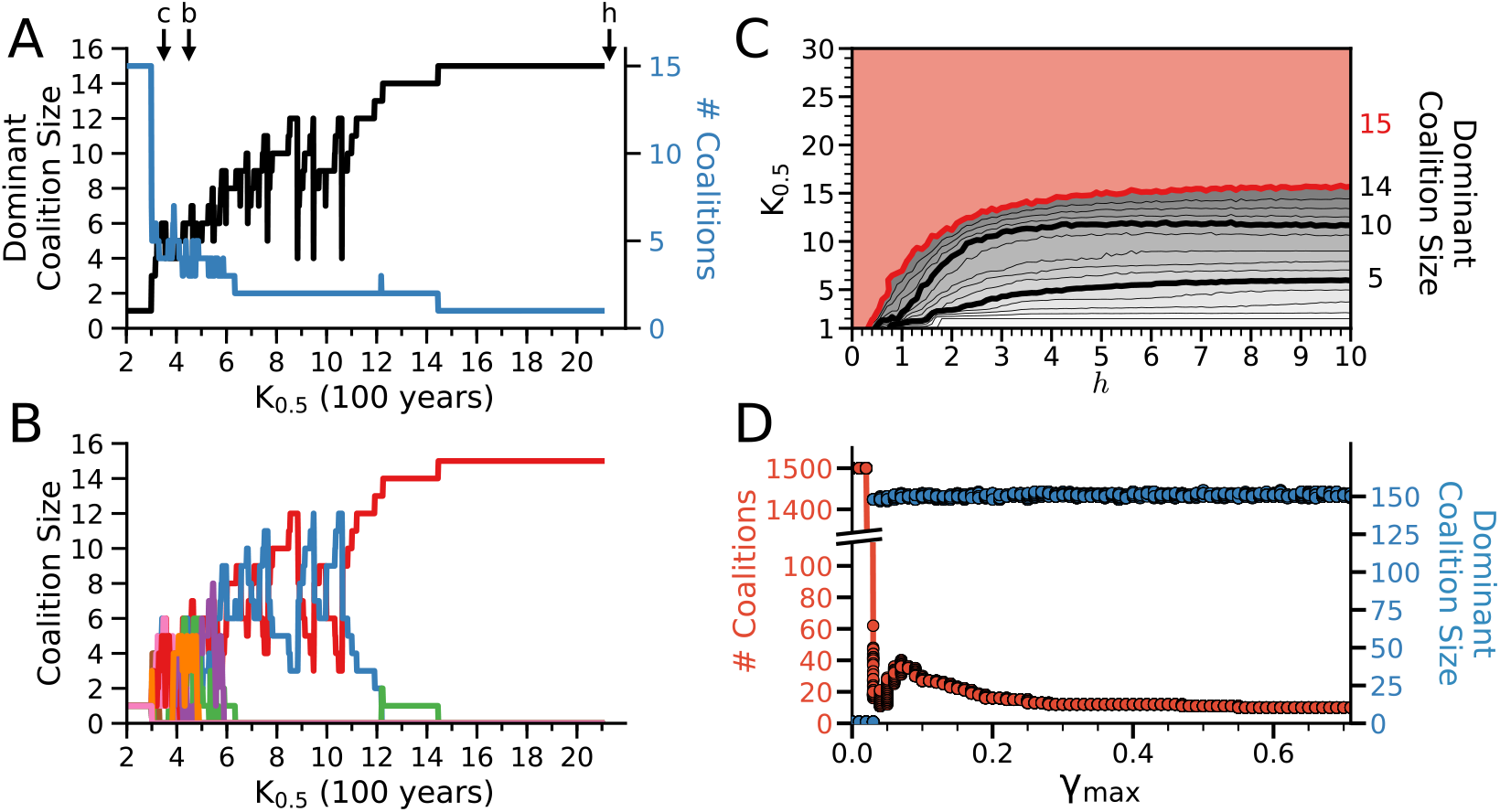
Effects of K_0.5_ and the Hill coefficient *h*. A. When the K_0.5_ value was increased every 100 years from a starting value of 1 to 20 in a population of 15 individuals with γ_max_=1.0 and *h*=2, the dominant coalition size (black) tracked the K_0.5_ value while the number of coalitions decreased (blue). Appropriate K_0.5_ values for gorillas (*g*), chimpanzees (*c*), bonobos (*b*) and simple forager humans (*h*) are indicated by arrows (see text). B. Corresponding evolution of coalitions as the K_0.5_ value increased. Coalitions with low DS perished due to defections into higher DS coalitions. Eventually all 15 individuals coalesced into single dominant coalition as K_0.5_ approached 15. C. Effect of *h* on dominant coalition size as the K_0.5_ was increased from 1 to 30, with γ_max_=1.0. Red area indicates K_0.5_-*h* combinations in which an egalitarian coalition of all 15 individuals was dominant, which occurred at lower K_0.5_ values for lower values of *h*. Data points are the median of 100 randomized trials of initial α and β values, for K_0.5_ and *h* incremented in units of 1 and 0.1 respectively. D. Effect of increasing γ_max_ on the dominant coalition size for a population of 1,500 individuals with K_0.5_=150 and *h*=6, showing a sudden transition from an alpha hierarchy to multiple co-dominant coalitions averaging approximately 150 members each for γ_max_ >0.03.

The effect of Hill coefficient *h* is illustrated in Fig. 5C in a population of 15 individuals with γ_max_=1.0. For high values of *h*, the dominant coalition size tracked the K_0.5_ value. For lower values of *h*, the dominant coalition size was larger than the K_0.5_ value. At *h*<0.5, the dominant coalition size was always the full population of 15 individuals regardless of the K_0.5_ value. This reflects the fact that for high values of *h* in Eq.(2), the value of gossip factor γ(*N*) plummets rapidly once the coalition size *N* exceeds the K_0.5_ value, at a faster rate than the sum 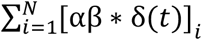 increases. For lower *h*, however, γ(*N*) falls less rapidly than the sum increases, so that coalition sizes larger than the K_0.5_ value have the highest DS.

Thus, the K_0.5_ and *h* values together determine dominant coalition size in a given population. For example, with γ_max_=1.0 and K_0.5_=150, the population remained fully egalitarian until the population size exceeded 150. When increased further to 1,500 individuals, population coalesced into (1500/K_0.5_)≈10 co-dominant coalitions averaging 150 individuals (range 149-151) in size when *h* was high (>6, Fig. 5D), and into a fewer number of larger co-coalitions as *h* was lowered until a single egalitarian coalition remained.

### A model prediction: weapon use and the egalitarian transition

Our model can be applied to make a prediction about the role of weapons in initiating the egalitarian transition, as has been hypothesized in coalitional enforcement theory [12]. In our model, the sole motivation for all individuals in the population is to dominate and avoid being dominated, i.e. to improve dominance rank by increasing their DS. Under these conditions, it is reasonable to assume that if weapons became available, the first instinct would be for individuals to use them to improve their dominance rank within the hierarchy. To simulate how a pre-egalitarian alpha hierarchy of primates would respond to the introduction of weapons, we modified our model to allow population size to vary by incorporating independent birth and death rates. The simulation in Fig. 6 illustrates a dominance hierarchy of 30 individuals with the gossip parameter γ_max_ at 0.03 (below the threshold for the egalitarian transition) and *h*=2.0, intended to emulate a chimpanzee community with 15 males and 15 females of which 10 are fertile. Based on ethnographic data [25, 26], the birth rate was set at 0.2 births/fertile female/year=0.067/individual/year, which was balanced by the same total death rate to achieve a stable equilibrium. The total death rate consisted of within-group murders (0.002/individual/year, based on a reported chimpanzee intragroup murder rate of roughly 200/100,000 [26]), death upon reaching age 50 (0.02/individual/year) and other deaths (infant mortality, accidents, infection, cancer, between-group murders, etc. totaling 0.045/individual/year). Under these conditions, population size varied stably between 26-31 individuals over a simulated 500 year period (Fig. 6). To simulate the effects of introducing weapons into this alpha hierarchical population, we used published data reporting that the overall within-group murder rate is similar among weaponless chimpanzees and weaponized human simple foragers (about 200/100,000), whereas the incidence of within-group violent physical attacks is 200-300 times higher among chimpanzees [26]. If we interpret this data to imply hypothetically that the lethality of physical attacks might increase 200-fold if pre-egalitarian primates in our model were to evolve weapon use, the resulting increase the mortality associated with within-group conflicts increases from 0.002 to 0.400/individual/year, increasing total mortality rate from 0.067/individual/year to 0.465/individual/year. Fig. 6 shows that when this higher mortality rate is introduced at 500 years into the simulation, the death rate outstrips the birth rate and the population becomes extinct within 90 years. This model prediction is consistent with the observation that chimpanzees and other non-human primates, although generally adept at tool use, have not evolved the ability to use tools as weapons [27, 28]. In contrast, once γ_max_ has exceeded the critical value for the egalitarian transition in our model, no individual can improve their DS or dominance rank by eliminating another individual, since the egalitarian coalition already has the highest DS, which will decrease if members are eliminated. Together, these model predictions suggest that the egalitarian transition may be a pre-condition for adaptation of tools as weapons (see Discussion).

**Fig. 6.**
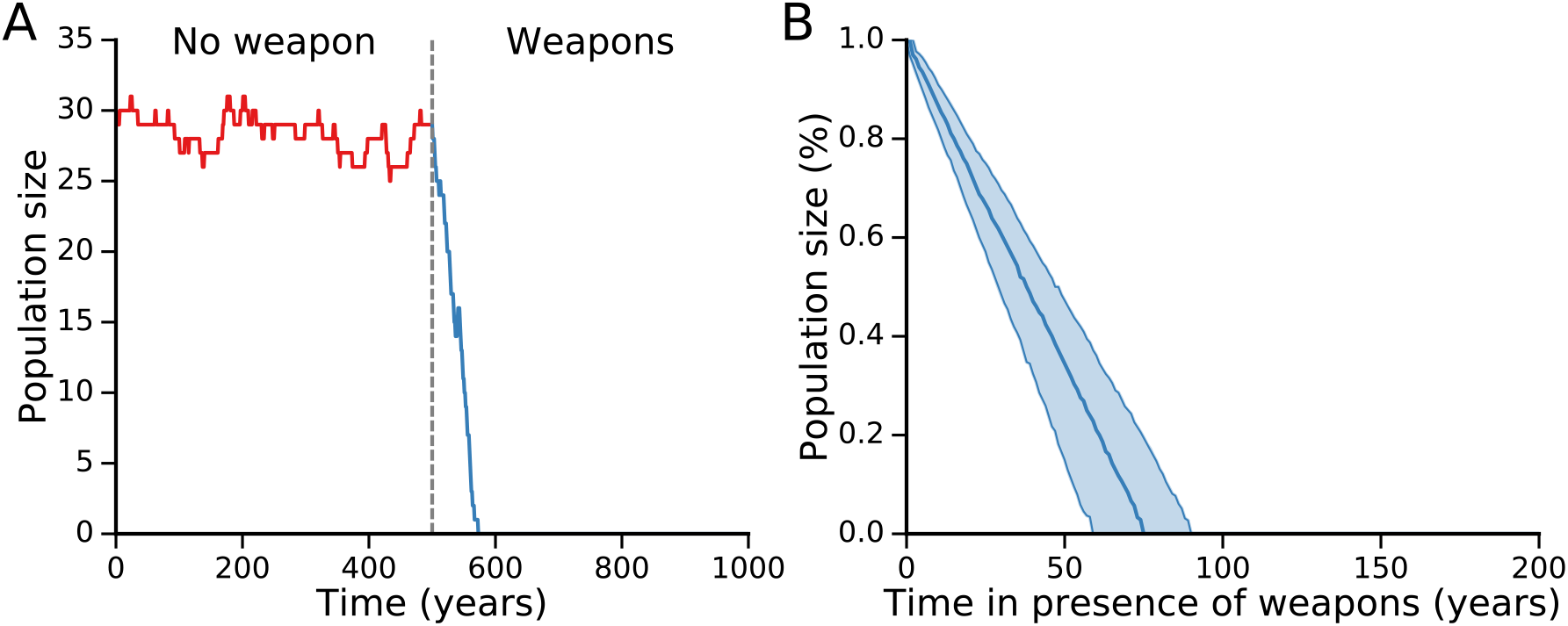
The effects of introducing weapons into an alpha hierarchy population. **A.** We simulated a chimpanzee community of 30 individuals (15 females of which 10 are fertile). Rather than automatically replacing individuals reaching the age of 50 with new individuals, the model was modified to incorporate independent birth and death rates, using data extracted from studies comparing wild chimpanzee and simple human forager populations [25, 26] summarized in the text. With model parameters corresponding to an alpha dominance hierarchy (γ_max_=0.03, *h*=2.0), the chimpanzee population size before the introduction of weapons remained stable for 500 years. At 500 years, the introduction of weapons (dashed line) was simulated by increasing the mortality associated with physical attacks by a factor of 200 (from 0.002 to 0.400/individual/year) as described in the text [26]. The resulting increase in total death rate (to 0.465/individual/year) outstripped the birth rate (0.067/individual/year), leading to extinction of the population within 90 years. **B.** For 1,000 different randomized initial values of α and β evolved for 500 years as in A, the medium time to extinction after introducing weapons was 75 years. Shaded areas indicate 95% confidence intervals [59 years, 90 years]. The y-axis is normalized to the population size after 500 years, just before the introduction of weapons.

## Discussion

The high prevalence of egalitarianism among human simple foragers living in small scale hunter-gatherer bands contrasts sharply with dominance hierarchies that characterize non-human primate societies, including great apes such as chimpanzees and bonobos presumed to be our closest relatives [3, 4]. Here we have developed a simple population model testing the plausibility that the invention of complex language (the gossip factor γ) could act as the critical parameter initiating an abrupt transition from a hierarchal to a fully egalitarian society, equivalent to a phase transition (Fig. 7). Unlike game theoretic models, our model does not include altruistic behavior as a strategy, yet the egalitarian transition with large-scale cooperation emerges nevertheless, as discussed below.

**Fig. 7.**
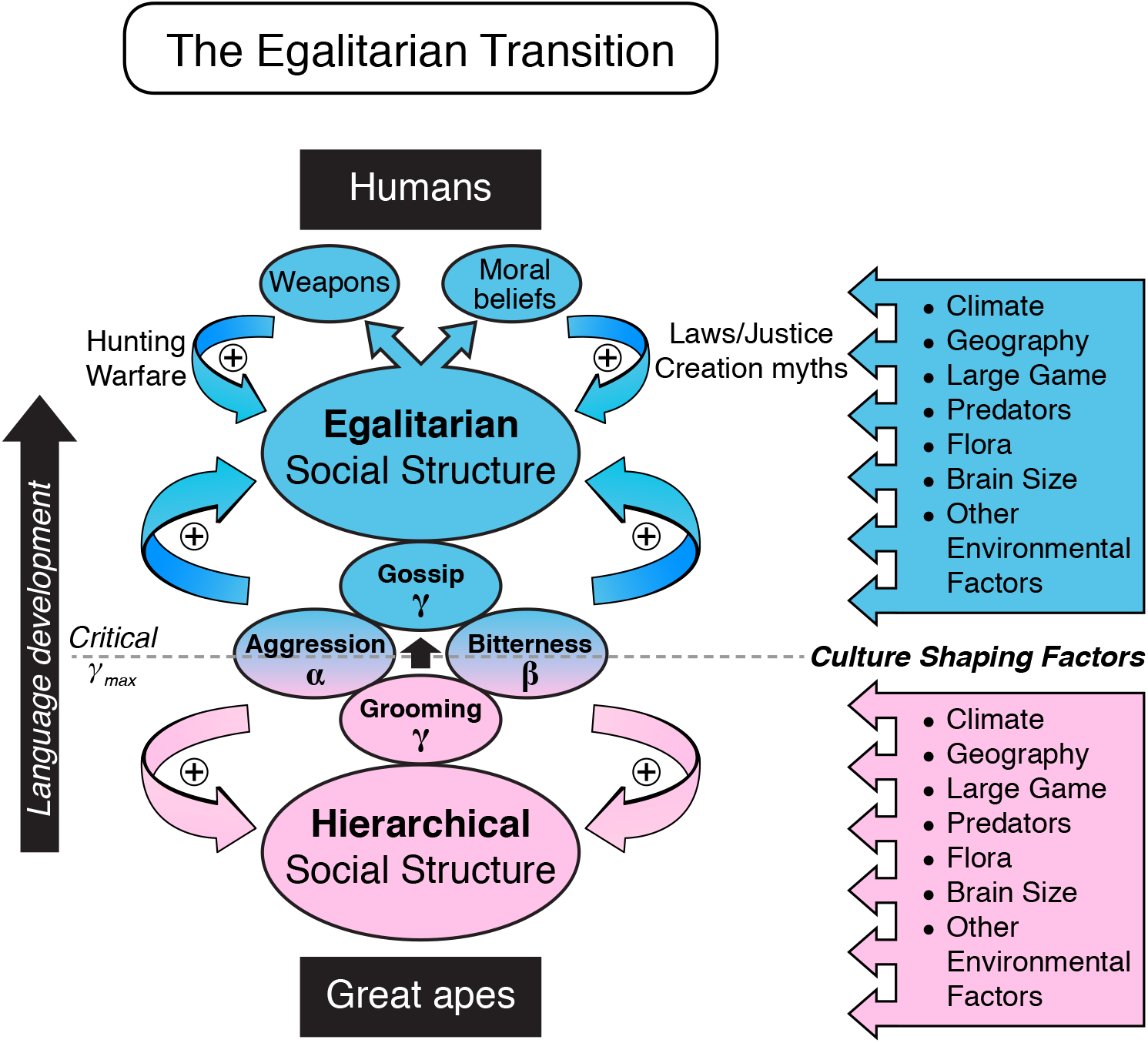
Factors underlying the egalitarian transition as language develops in early humans. In great apes, grooming (low γ) combines with aggressiveness (α) and bitterness (β) to cements a hierarchical social structure; in early humans, gossip (high γ) replaces grooming and destabilizes the individual alpha position by allowing large coalitions to form, causing a sudden transition to egalitarianism at a critical value of γ_max_. Once the egalitarian transition has occurred, it is further strengthened by development of weapons and moral beliefs promoting altruistic behavior. The egalitarian transition is also determined by local culture-shaping factors that influence α, β and γ.

### Model features

Our key assumption is that stochastic gossip-driven interactions will encourage individuals to join coalitions that improve their DS so as to optimize their dominance rank. For simplicity, we avoided the step in evolutionary game theory paradigms that explicitly evaluates the costs and benefits of various actions in controlling an individual's decisions. Instead, these costs and benefits are folded into the DS, with higher DS value implicitly reflecting a higher benefit-to-cost ratio of joining a coalition. The model algorithm then stochastically explores different possible coalitionary combinations until an optimal power-sharing configuration is reached in which no further rearrangements can improve the DS of any individual. This is analogous to a thermodynamical energy landscape in which random heat-related motion causes the system to explore new configurations until it finds a stable energy well from which it has a low probability of being dislodged by random events. In the case of melting ice, for example, individual water molecules in a highly ordered three-dimensional array cemented together by intermolecular hydrogen bonds are overcome by heat-driven vibrational forces causing a transition to an amorphous fluid at a critical temperature. In our model, individuals in a highly ordered one dimensional array of a dominance hierarchy determined by their aggressiveness-bitterness factor ±² (analogous to the energy stored in intermolecular hydrogen bonds between adjacent water molecules) are destabilized by gossip (γ)-driven energy (analogous to heat causing vibrational kinetic energy of water molecules). Thermodynamically, heat must be added to break down the structure of ice at 0°C during the phase transition to liquid water at 0°C (the heat of formation), corresponding to an increase in entropy. In our model, gossip, like heat, breaks down the hierarchical structure into a fluid power-sharing egalitarian state, generating an increase in coalition entropy S_c_ (Fig. 4C). Egalitarianism emerges when the parameter γ_max_ reaches a critical threshold, at which point the dominant alpha position suddenly becomes unstable for any individual. At this threshold, any individual alpha can be toppled by an anti dominance coalition of resentful lower-ranking individuals. Thus, once the gossip parameter γ_max_ exceeds its critical value, it is in the direct self-interest of each individual, including the alpha, to avoid dominant alpha behavior, knowing that the inevitable consequence will be a precipitous fall in sociopolitical rank. Thus, pure self-interest motivates each individual to join a coalition, the direct benefit of which is an increase in their DS and dominance rank, and the implicit cost of which is to share risk of injury, power and resources equitably with other members of the coalition. Notably, the cost per coalition member of injury when confronting an alpha decreases as coalition size and DS increases relative to that of the alpha, driving larger and larger coalition sizes until the full population becomes egalitarian (assuming *N* < K_0.5_). Since the alpha at the top is universally resented and so has no natural allies, the alpha has no choice but to join the coalition or be relegated to the lowest dominance rank. This phase transition from hierarchy to egalitarianism at a critical threshold of language development has parallels in studies of human cooperation, in which phase transitions from defectors to cooperators occur when critical penalty thresholds are reached that punish free-riders in the public goods game [18, 19]. Importantly, our model remains robust when free-riders are included by adding the γparameter (Fig. 3). Moreover, the critical γ_max_ value corresponding to the transition from alpha hierarchy to egalitarianism could be derived analytically (Eq.(4)), as validated by excellent agreement with numerical simulations for cases in which the K_0.5_ and *h* values promoted an egalitarian transition of the full population (*N* < K_0.5_ or low *h*). Thermodynamics-inspired approaches have also previously been proposed in economics [29] and informatics [30].

An important feature of our model is that it does not preclude non-dominant coalitions that create a hierarchy of coalitions within coalitions (as encompassed by the coalition entropy calculation shown in Fig. 4). However, all non-dominant coalitions remain subservient to either the alpha individual or the full egalitarian coalition depending on the γ_max_ value. Thus, the model contains an inherent flexibility that can accommodate many local power-sharing non-dominant coalitions under the umbrella of a globally egalitarian society.

The Hill equation formulation in the model limits the maximum size of a dominant coalition by decreasing the benefit/cost ratio (i.e. reducing the DS) of forming/maintaining a coalition beyond a critical size determined by K_0.5_ and *h*. Egalitarianism requires individuals to gossip frequently enough to be able to cement friendships and judge each other's intentions and trustworthiness, which is key to reducing the costs of coalitional enforcement. The time utilized for gossip, however, must be balanced against other needs (hunting, food gathering, sleeping, etc.), limiting the maximum coalition size before egalitarian decision-making becomes impractical. This maximum coalition size has been estimated to be near 150 in modern humans [20]. In our model, setting the K_0.5_ value to 150 with a sufficiently steep Hill coefficient naturally mimicked this situation (Fig. 5D), with the implication that beyond this coalition size, gossip-based egalitarianism becomes impractical. In this case, the population self-organizes into competing co-dominant coalitions, which, when resources are limited, potentially leads to warfare, expansion into new territory, or re-emergence of hierarchical structures in the form of big men, chiefs, etc. to represent the interests of competing co-dominant coalitions. There are examples of larger human groups which maintain an egalitarian ethos, such as the Iroquois Confederation [31] and 18^th^ century Montenegrin tribal societies [32]. Our model can incorporate these cases by increasing the K_0.5_ value or decreasing *h* value to reflect underlying cultural factors that allow egalitarian societies >150 in size. Thus, the K_0.5_ and the Hill coefficient *h* values reflect the composite effects of communication capabilities, psychological temperament and cultural factors (transition from nomadic to sedentary life styles, agriculture and domestication of animals engendering property ownership issues, etc.) whose interplay determines the maximum size of dominant coalitions as human societies become larger [4, 7, 33, 34].

The Hill equation also allows the model to be adapted to represent coalition development in nonhuman primates if g is generalized to include non-verbal forms of communication, such as grooming [20]. For example, in chimpanzee communities, whose size can average near 50 individuals with approximately 15 adult males eligible for the alpha position, the alpha position is often shared by a coalition of several co-dominant males [35]. In our model this would correspond to a superthreshold value of γ_max_ and K_0.5_=2-3 (Fig. 5A, arrow labeled *c*). Bonobos, on the other hand, have similar sized communities as chimpanzees but have evolved a matriarchal power structure in which small female coalitions or mother-son coalitions dominate individual males [28]. Our model could account for this power structure if male bonobo temperament is not suitable to form stable male-male coalitions, such that γ_max_ remains subthreshold, but a more suitable female bonobo temperament is compatible with a superthreshold γ_max_ and a K_0.5_ value of 2 or greater, such that the coalition DS of multi-female and mother-son dyads can exceed that of individual males (Fig. 5A, arrow labeled *b*).

Finally, our model makes an important prediction that may explain why non-human primates in the wild, among whom tool use is common, do not use tools in a directed manner as weapons [27, 28]. When we simulated the introduction of weapons into a hierarchical chimpanzee community of 50 individuals, such that the mortality rate of physically violent challenges became equivalent to that in weaponized human simple human foragers (i.e. 200-300 times greater), the total death rate outstripped the birth rate [25] and the population became extinct in <100 years (Fig. 6). The implication from our model is that the egalitarian transition may have been a cultural precondition for the adaptation of tools into weapons (and therefore also a precondition for large game hunting and weapon-based warfare). That is, until egalitarianism created social sanctions to prevent individuals from using weapons to improve their within-group dominance rank and instead restricted weapon use to hunting and between-group warfare, their introduction into a hierarchical community would have led to rapid extinction. Once egalitarianism has become established, however, the introduction of weapons facilitating killing-at-a-distance markedly reduces the cost of coalition enforcement, as hypothesized by Bingham [12], with the immediate effect of stabilizing and locking in place the egalitarian structure. Moreover, weaponized egalitarian bands would then gain a tremendous fitness advantage over competing hierarchical bands. Advances in weapons technology could subsequently drive consolidation of human simple forager bands into larger and larger groups, as posited by coalitional enforcement theory [12].

### Comparison to previous coalition theory models

The vast majority of theoretical models investigating coalition formation have been game theoretic approaches predicting coalitionary strategies that maximize individual fitness depending on the costs and benefits of different actions and the information available to the individuals [14, 36]. The actions typically include selfish (win-lose), altruistic (lose-win), mutually beneficial (win-win), or spiteful (lose-lose) choices. Despite the obvious advantages of within-group cooperation when groups compete for common resources, however, how groups become initially enriched in cooperative individuals is still highly debated [21-24]. That is, if only the fittest survive, why should one perform an altruistic act that is costly but benefits another, or contribute to the public good if free-riders can enjoy the same benefits for free? In other words, since cooperation is costly, within-group natural selection favors free-riders over altruists, especially when group size exceeds more than a few individuals [5, 36]. Large-scale cooperation is facilitated when punishment and/or reward are added as options, although evolutionarily stable cooperative strategies [37] immune to corruption by free-riders are still not generally robust [5, 18, 19, 36, 38-40]. In this context, punishment is considered to be more effective than reward at stabilizing large-scale cooperation by protecting altruists from exploitation [19, 41, 42]. Our simple model is not only generally consistent with these findings, but by showing that the egalitarianism transition can be driven purely by self-centered motivations (will to dominate and resentment against being dominated), may even provide some insight into how large-scale cooperation can arise in the absence of genuine altruism. In this case, the incentive for cooperating is not altruistic, but rather the fear of being punished for not cooperating.

The phase transition to egalitarianism in our model as γ_max_ increases also has counterparts in evolutionary game theory models of human cooperation. For example, the public goods game commonly exhibits phase transitions in which the population suddenly shifts from defectors to cooperators in response to small changes in punishment and reward parameters [18, 19]. In our model, this corresponds to the point at which the cost of punishment by the coalition exceeds the benefit of remaining outside of the coalition. Pandit et al. [39] found that the combination of low coalition costs and equitable male access to females for reproduction could lead to large scale coalitions, which they speculated may be relevant to egalitarianism in human simple foragers. The speculation about lower coalition costs is consistent with our model's formulation, since the DS increases as coalition size increases up to K_0.5_, implying that the implicit coalition costs are lower and benefits are greater as coalition size increases.

As an alternative to game theoretic approaches, Gavrilets et al [17] developed a coalitional formation model based on the theory of dynamic linking and network formation, in which explicit evaluation of costs and benefits of certain actions in controlling decisions were replaced with an affinity matrix defining each individual's affinity to assist or not assist other individuals during a conflict, that evolved with experience. Relevant to our model as well, the authors argue that *“our approach is justified not only by its mathematical simplicity but by biological realism as well. Indeed, solving the cost-benefit optimization tasks (which require rather sophisticated algebra in modern game theoretic models) would be very difficult for apes and early humans especially given the multiplicity of behavioral choices and the dynamic nature of coalitions. Therefore treating coalitions and alliances in early human groups as an emergent property rather than an optimization task solution appears to be a much more realistic approach.”* Using this agent-based model, they also identified conditions in which the population underwent a sudden phase transition to full egalitarianism analogous to our model. The random assignment of affinities in their model, however, had no explicit relationship to language development or other factors proposed to be important in promoting the egalitarian transition in early humans. Our model, on the other hand, is conceptually similar in terms of avoiding an explicit cost-benefit formulation, but is simpler and directly links the egalitarian transition to language development. Rather than randomly assigning affinities, our model characterizes each individual with the parameter αβ reflecting the combination of their will to dominate (aggressiveness) and resentment against being dominated (bitterness), similar to resource holding potential (RHP) concept [36]. α² changes only with age, but otherwise is not altered by experience as in the agent-based affinities model. Language skills provide individuals with opportunities to advance their rank by joining coalitions with higher DS, who interact randomly until an optimal power structure is achieved at which point no individual can further improve their dominance rank by either remaining alone or joining a new coalition. This process naturally leads to a sudden phase transition from hierarchical to egalitarian power structure when γ_max_ reaches a critical value. Thus, we arrive at similar phase transition behaviors as both Gavrilets et al [17] and human cooperation game theory studies [18, 19] using a novel approach aligned with thermodynamical concepts.

### Model limitations

Like the Gavrilets et al [17] affinities network model, our simple model does not explicitly compute the evolution of the egalitarian transition in the classical sense that something evolves only if the benefits outweigh the costs, since we have not explicitly formulated costs and benefits of gossip or decisions to attempt or not to attempt to dominate. Rather, our model explores the dominance landscape of all possible coalitions to identify the optimal power-sharing arrangement which optimizes the DS for all individuals in the population. Our assumption is that through stochastic interactions, the system will eventually arrive at the most stable configuration, analogous to a thermodynamical process. It could be argued, however, that without explicitly simulating the system's evolution, there is no guarantee that the most stable configuration will be achieved or will be evolutionarily stable. In this sense, our model should be considered a description of a possible scenario that explains the egalitarian transition as a phase transition related to language development, rather than model of the evolution of the actual process.

Our model also makes no attempt to explain how or why complex language developed in alpha hierarchical primates prior to the egalitarian transition. Language may have evolved as a behavioral coordination tool to confer group protection against predators [15], but then gradually supplanted grooming as the major form of socialization among humanoid primates. We simply assume that once complex language permitting gossip developed, it promoted large coalition formation by allowing third party information to be communicated efficiently throughout the coalition. In the absence of complex language, it is unclear how primates could negotiate or enforce large scale coalition agreements [8, 14, 15]. As stated by David-Barrett et al. [15], for example, *“Maintaining coordination in such large groups requires both a large brain and the ability to communicate third-party information. Third-party information, however, cannot be passed on without some form of language because this requires both time and place to be marked as well as the ability to identify third parties and comment on their action.”* Thus, we argue that by affording the capability to express abstract concepts as well as emotions to third-parties, complex language is inherently superior to grooming or simple language for promoting and maintaining large coalitions as well as developing proactive strategies to punish individuals exhibiting alpha behavior [8, 14, 15]. Complex language facilitates the formulation of collectively agreed upon transmissible rules of acceptable and unacceptable behavior punishable by cultural status leveling mechanisms such as ridicule and shaming to limit escalation of conflicts to lethal violence, especially after the adaptation of tools as weapons markedly increases the potential lethality of physically violent challenges.

Our model assumes that the dual desires to dominate and to avoid being dominated encompassed in the aggressive-bitterness factor α² serve as sufficient motivation for an individual to join a coalition if it improves their dominance rank and thereby gives them greater access to reproductive and other resources. We arbitrarily assumed that time scale for coalition formation is rapid, i.e. within a year, and once formed, coalitions persist for at least a year, after which the coalition DS is recalculated. This is a simplification, since coalitions may form and persist or disappear at different rates.

It might be argued that nothing in our model specifies that the sociopolitical structure within a coalition is egalitarian. However, we argue that although our model does not dynamically evolve resource division or decisions supporting collective actions or free-riding according to prespecified rules as in game theoretic approaches, the implied sociopolitical structure of coalitions in our model is inherently egalitarian, since: i) resentment against being dominated by others is the major driver of coalition formation; ii) the individual DS of all members of a valid coalition is less than the coalition's collective DS, so each member is better off as a coalition member and no member can exhibit alpha dominance behavior without provoking resentment and retaliation by the coalition. Moreover, the only specific requirement for gossip is to allow a group of individuals to communicate effectively enough to agree to come to each others* aid should a bully try to dominate a coalition member, analogous to genetically-encoded coalition formation by which primates and other mammals instinctively defend their offspring against aggressors.

To incorporate free-rider tendencies into coalitions in our model, each potential coalition's DS was multiplied by the laziness factor γ(a randomly selected value from [0,1]) to reflect, in a random fashion, contamination by free-riders. Thus, the laziness factor γstatistically summarizes how individual dynamics within a coalition reduce its overall DS (i.e. fitness) while avoiding the need to compute the costs and benefits of free-riding decisions explicitly. A limitation is that this approach precludes the ability to assess whether the optimized power-sharing structure is an evolutionarily stable strategy by testing its robustness to invasion by alternative strategies [37]. However, our model implicitly allows only selfish (win-lose) and punishment (lose-lose) choices, motivated by aggressiveness and bitterness, respectively. Since the final optimized power-sharing state is defined as a configuration in which no further defections can improve any individual's dominance ranking, we argue that if our model were converted to a game theoretic formulation, the final state is likely to be evolutionarily stable with respect to selfish and punishment choices. However, it may be evolutionarily susceptible to invasion by altruistic (lose-win) or mutually beneficial (win-win) choices, which in game theory models can generate populations that oscillate between periods when cooperators gain the upper hand and periods when free-riders dominate [5, 36, 38, 40]. This could be an important factor promoting consolidation of competing bands into larger human populations (such as simulated in Fig. 5D with 10 coalitions of approximately 150 individuals each). That is, assuming that oscillations are out-of-phase in different coalitions, coalitions in the cooperative phase would have an advantage in warfare over coalitions in the uncooperative phase, driving consolidation of human populations through conquest, as proposed previously [11, 12, 43].

Our model takes a high altitude view of primate societies, without an attempt to account for local socio-ecological and environmental factors that may play key roles in shaping the culture of individual societies, including the degree of egalitarian versus hierarchical sociopolitical structure [2, 10]. However, the observation that great apes uniformly exhibit dominance hierarchies, whereas small-scale simple forager human societies studied across multiple continents and diverse socio-ecological conditions commonly exhibit egalitarian societies [4] suggests that a universal mechanism(s), largely independent of local environmental factors, may be at play. Local factors, on the other hand, are likely to significantly influence parameter values in our model to synergize or antagonize the egalitarian transition (Fig. 7).

The Hill equation is a purely heuristic way to model the effect of group size on the ability to maintain effective coalitions, but serves the purpose in the model of restricting the maximum size of fully egalitarian societies to approximately the K_0.5_ value, thus accounting for the common re-emergence of hierarchical structures when human populations exceed this size. For ancestral humans in which complex language evolved to replace grooming as the preferred method of social interaction [20], the K_0.5_ value would be limited by the number of individuals with whom it is practical to gossip frequently enough to reinforce sociopolitical ties, which has been estimated to be around 150 in modern humans [20]. Thus, for human simple foragers whose band size is typically much less than 150, the population would immediately transition to full egalitarianism, as predicted in Fig. 2. Even if the K_0.5_ value were ten-fold lower in early humans, egalitarianism would still emerge when γ_max_ achieved a critical value.

Finally, we choose to simulate a band of male hunter-dominated simple foragers, in which females typically have a less obvious power-sharing status [1, 4]. However, the individuals in our model could equally well be taken to represent male-female couples, especially in monogamous simple foragers, in which the aggressiveness-bitterness factor α² is strongly influenced by the natural inclination of male-female couples to motivate and protect each other.

### Origins of moral beliefs and weapon use

In addition to promoting large scale coalition formation through gossip, language development provides the means to formally codify acceptable and unacceptable behaviors into a moral belief system which can be communicated intergenerationally throughout the group, and serve as the basis for developing cultural status leveling traditions (ignoring, ridiculing, shunning, exiling and even executing free-riders and bullies) that are a universal feature of human simple forager bands [4]. A moral system based on fear of punishment by the egalitarian coalition establishes vigilance against free-riders and bullies so that genuine behavioral and/or psychological altruism can subsequently strengthen the cooperative one-for-all, all-for-one egalitarian ideology. Thus, the Golden Rule philosophy that is ubiquitous among extant simple forager egalitarian bands would in our model begin as *“Do unto others as you would have done unto yourself, or else you will be punished”* and then subsequently evolve into *“Do unto others as you would have done unto yourself, because it is the right thing to do (and if not, you will be punished)”* encompassing both punishment-based and reward-based elements. As noted above, this outcome has been generated by agent-based game theoretic models in which altruistic traits are stabilized if a high fraction of the group serve as “punishers” who increase the cost ratio for antisocial free-riders and bullies [5, 36, 38], or as “rewarders” who encourage prosocial altruistic acts [40]. Since the moral belief system emerges as a collective consciousness (or superego) of the group, it is not unreasonable to speculate that the human imagination might attribute moral rules to supernatural forces in the form of a creation myths and deities, which further reinforce the egalitarian credo (Fig. 7).

## Materials and Methods

All simulations were performed on a personal computer using custom-written software in the Julia programming language [44, 45]. To obtain a truncated exponential distribution, values were drawn using the formula 0.05*10^RN^ where RN is a random number between −2 and 0, yielding a range from 0.005 to 0.5 with a median of 0.05. Gaussian and random distributions ranging from 0.005 to 0.5 were obtained from the Distributions.jl julia library. Conventional percentile bootstrap-resampling approach with 10,000 replications was used for estimating 95% CIs [46–48].

## Acknowledgments

We thank Christopher Ko, PhD and Zhilin Qu, PhD. for their helpful contributions.

